# Longitudinal analysis of glioblastoma response to oncolytic HSV1 guides PLX3397 combination for enhanced viral permissiveness

**DOI:** 10.1101/2024.09.17.613527

**Authors:** Hirotaka Ito, Naoyuki Shono, Claudia Zagami, Min J Kim, Alayo A. Quazim, Eric McLaughlin, William F. Goins, Soledad Fernandez, E. Antonio Chiocca, Hiroshi Nakashima

## Abstract

Clinical trial data underscores the need to improve oncolytic virus (OV) distribution within tumors, a challenge compounded by the lack of predictive biomarkers and limited opportunities for post-treatment analysis. To decipher the factors influencing treatment outcomes, we employed multimodal bioluminescence imaging (MM-BLI) in conjunction with magnetic resonance imaging (MRI) to monitor OV infection, replication, and tumor viability in orthotopic brain glioma mouse models. This approach revealed two distinct therapeutic responses: “Responders” with tumor regression and “Non-Responders” with tumor progression. In-depth analysis of individual brains from both groups uncovered dynamic interactions between the OV and the tumor microenvironment, highlighting the involvement of Iba-1+ microglia and tumor necrosis in hindering OV distribution within the tumor. To address this, we incorporated a CSF-1 receptor inhibitor (PLX3397), which improved tumor control by enhancing OV’s direct cytopathic effects and reducing microglial interference. Our findings highlight microglia as a significant barrier to effective OV therapy, suggesting that targeting brain-resident immune cells could enhance the therapeutic efficacy of OVs in resistant brain tumors.

## Introduction

Oncolytic viruses (OVs) are engineered to replicate selectively and be toxic to cancer cells^1–6^. They replicate within tumors, spreading the infection through layers of tumor cells. This process induces immunogenic cell death, activating innate inflammatory responses by interferons and immune cells such as NK cells and mononuclear cells. While these responses eliminate the OV, they also trigger adaptive immune responses (CD4+ and CD8+ T cells), leading to potential long-term immunity against the tumor.

However, achieving effective distribution of injected OV within tumors, especially in cases like glioblastoma (GBM), remains challenging. Clinical trial data indicates limited dispersion and rapid clearance of injected OV in GBM tumors^7–10^. Furthermore, the varying therapeutic responses observed in preclinical studies utilizing immunodeficient mice with human GBM orthotopic xenografts emphasize the need to understand the interplay between direct cytotoxicity and the early innate immune response following intratumoral OV inoculation. Quantitative analyses of OV-tumor kinetics have been limited in previous studies, often focusing on viral activities, tumor burdens, or short timeframes due to the absence of robust biomarkers for simultaneous and long-term analysis of OV-tumor kinetics *in vivo*. To address this limitation, we integrated quantitative analyses with immunohistochemistry and transcriptomics, performed on individual mice^11–17^. This comprehensive approach allowed us to identify key factors influencing the therapeutic efficacy of OV therapy, offering valuable insights for the successful translation of OV therapy for brain tumors.

In our ongoing efforts to monitor the replication kinetics of oncolytic herpes simplex virus (oHSV) within tumors, we previously integrated two luciferase cDNAs separately into the oHSV genome^17^. These luciferases, *Renilla* luciferase (Rluc) and *Firefly* luciferase (Fluc), controlled by the HSV immediate-early (IE) promoter (as an infection marker) and the HSV late (L) promoter (as a replication marker), respectively, facilitated tracking of viral activities. In this study, we have advanced our approach by engineering the oHSV to simultaneously monitor both IE and L promoter activities. Furthermore, we engineered human GBM cells to express a secreted luciferase from *Cypridina* (Cluc), which is readily assayed in cultured medium and mouse serum^18, 19^. This intense imaging approach, termed Multimodal Bioluminescence Imaging (MM-BLI), allows for *in vivo* real-time longitudinal assays of both oHSV infective and replicative kinetics and mouse GBM tumor viability during virotherapy. Further, we show that these serial imaging modalities can be employed to validate agents that could be used to increase oHSV biodistribution in tumors, leading to the discovery of colony stimulating factor (CSF)-1 receptor antagonist, PLX3397, that significantly improves oHSV biodistribution in injected human GBM xenografts in immunodeficient mice.

## Results

### Engineering of luciferase-expressing oHSV and GBM cells

We conducted a comprehensive assessment of the longitudinal kinetics of oncolytic herpes simplex virus (oHSV) activities, aiming to understand their influence on tumor lytic efficacy and progression at advanced stages. New MM-BLI armed oHSV, iNG34^20^, is a NG34-based oHSV, incorporating dual luciferase genes controlled by HSV promoter cassettes activated during the immediate early (IE) or late (L) stages of the viral replication cycle (**Extended Data Fig. 1a**). We observed both luciferase activities exhibited temporal increases over time in iNG34-infected Vero cells. IE-promoter driven Rluc activity preceded the L-promoter driven Fluc, reflecting distinct kinetics of IE promoter activation compared to the L promoter under phosphonoacetic acid (PAA), which inhibits viral DNA synthesis (**Extended Data Fig. 1b, c**). Temperature-shifted assay at a non-permissive condition resulted in a significant reduction in Fluc activity (**Extended Data Fig. 1d, e**). In contrast, IE promoter-driven Rluc activity displayed smaller changes under these conditions, consistent with our earlier report^21^. These results support that the temporal kinetics of the viral lytic cycle are effectively measured using these dual luciferase biomarkers. Next, two human glioblastoma (GBM) cell line, namely U87ΔEGFR and GBM9 (G9), were also genetically modified to express *Cypridina* luciferase (Cluc), whose secretion is correlated with numbers of cultured cells (**Extended Data Fig. 2a**). This Cluc in sera obtained from peripheral blood increased after implantation of U87ΔEGFR-Cluc (**Extended Data Fig. 2b, c)** and G9-Cluc (**Extended Data Fig. 2d, e)** in mouse brains, and exhibited a robust correlation with T2-weighted tumor volumes (**Extended Data Fig. 2c, e)**, even though U87ΔEGFR-Cluc and G9-Cluc display distinct tumor formation, as the latter represents pseudopalisading necrosis (**Extended Data Fig. 2f-i**). These findings strongly indicate that serum Cluc is a readily accessible biomarker for monitoring the viability of these GBM cells in brains.

### Analyzing oHSV Activity and Tumor Viability In Longitudinal MM-BLI Study

We carried out a prolonged 60-day Kaplan-Meier (K-M) survival experiment concurrently with scheduled MM-BLI analyses following intratumoral iNG34 treatment on day 7 (as shown in (**Figure 1**). Consistent with prior reports^20, 21^, we observed a significant increase in the survival rate of mice bearing U87ΔEGFR-Cluc and G9-Cluc tumors when treated with iNG34 compared to the Vehicle control (**Figure 1a**). MM-BLI analysis distinguished patterns of luciferase activities when comparing long-term survivors (complete responders, CR), mice that initially showed signs of response but later succumbed (partial responders, PR), and mice that perished from their tumors despite iNG34 treatment (non-responders, NR) (**Extended Data Fig. 3 and Figure 1b-g**). We grouped CR and PR mice together as responders. In contrast to U87ΔEGFR-Cluc model, G9-Cluc tumor model represent little difference in NR group compared to vesicle group and lack CR group. As Cluc levels decreased, so did oHSV’s Rluc and Fluc activities. In these responder mice, we noted an initial peak in both oHSV Rluc and Fluc, accompanied by a swift decline in tumor Cluc levels, which essentially were undetectable by day 20 in U87ΔEGFR-Cluc model and day 10 in G9-Cluc model. In PR mice, Cluc levels started rising again after the decline of oHSV Rluc and Fluc, indicative of occurring recurrent. In Non-Responders, despite the initial peak in both oHSV’s Rluc and Fluc, tumor Cluc levels also increased. Notably in G9-Cluc model, both oHSV’s Rluc and Fluc reascended upon recurrence events (as inclining Cluc) in both NR and PR groups. These different trends observed in two GBM models suggests that G9-Cluc tumors is more oHSV-resistant than U87ΔEGFR-Cluc due to more complicated tumor environment (**Extended Data Fig. 2f-i**). Our initial studies in two orthotopic human GBM models demonstrate the advantage of integrating longitudinal MM-BLI with survival experiments, revealing that the dynamic changes between the oHSV and tumor are closely involved in the therapeutic responsiveness in oncolytic virotherapy.

**Figure 1.**
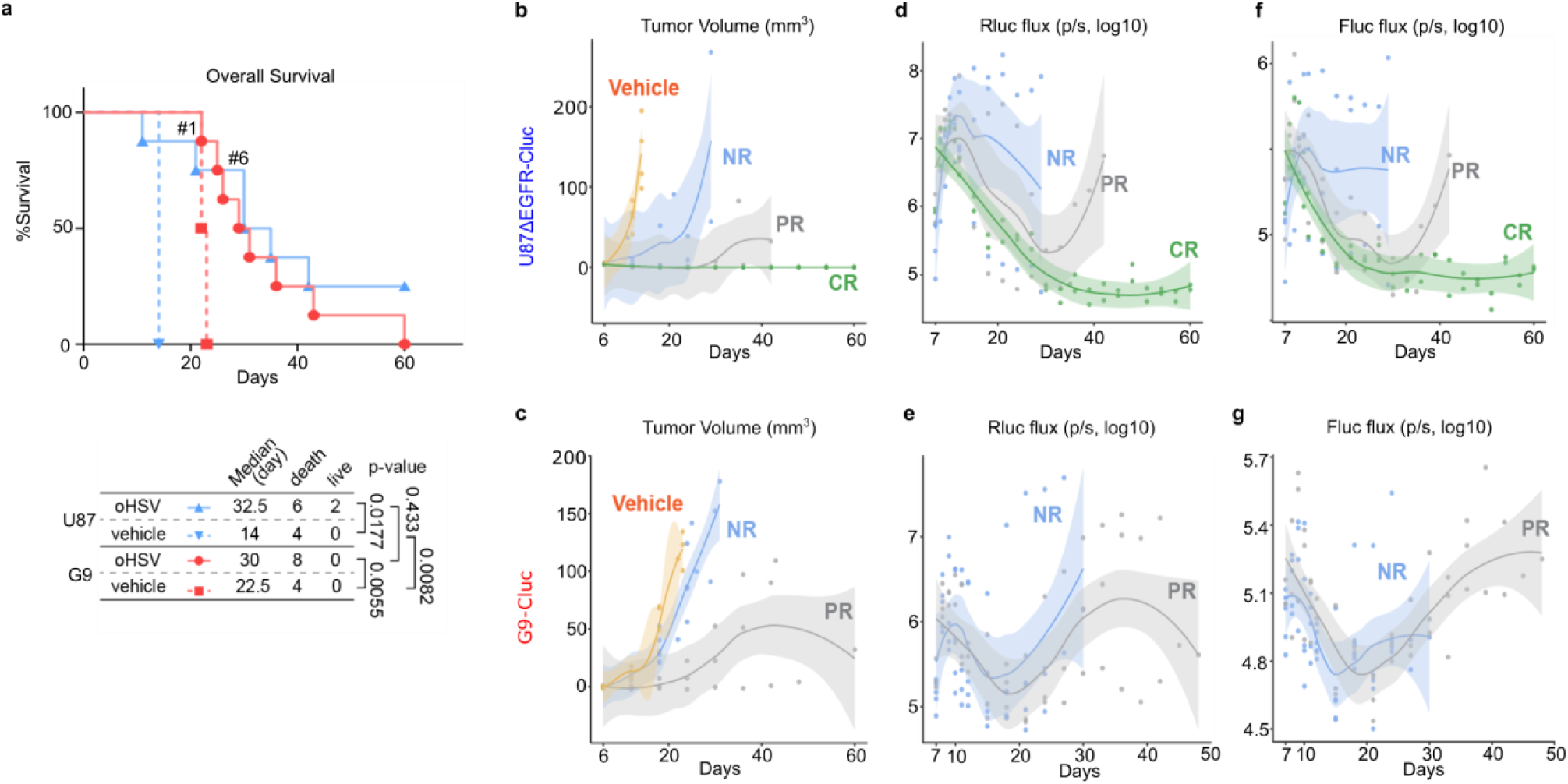
Multiplex temporal analysis in human glioblastoma intracranial (i.c.) xenografts after virus inoculation. A total of 100,000 U87ΔEGFR-Cluc or G9-Cluc cells were stereotactically implanted in the right cerebral hemisphere of female athymic mice. On day 7, 200,000 pfu of iNG34 (n = 8) or HBSS (n = 4) was inoculated at the tumor site. Blood was collected for Cluc measurement, and viral biomarkers (Rluc and Fluc) were assessed by IVIS for the first 5 consecutive days post-inoculation, followed by measurements every three days until the end of the experiment. **a,** Kaplan-Meier survival curves for U87ΔEGFR-Cluc (blue line) and G9-Cluc (red line) i.c. xenografts, with follow-up periods of 60 days after tumor implantation; P values are from the Log-rank test. Mice “#1” and “#6” indicate those that died on days 21 and 27, respectively, as shown in **Extended Data Figure 3**. **b, c,** Temporal analysis of tumor volumes calculated from serum Cluc levels, categorizing mice into Complete Responders (CR), Partial Responders (PR), or Non-Responders (NR) for U87ΔEGFR-Cluc, and PR or NR for G9-Cluc. **d-g,** Temporal analysis of viral biomarkers, Renilla luciferase (Rluc; **d, e**) and Firefly luciferase (Fluc; **f, g**). The y-axis represents flux (p/s, log10); scatter plots (dots), non-linear regression curves (solid lines), and two-sided 95% confidence intervals (shaded areas) are shown for each graph.

### Positive Correlation Between MRI T2 LIV and MM-BLI

Previous experiments using the U87ΔEGFR-Cluc model identified distinct patterns of iNG34 Rluc and Fluc expression between Responder (CR/PR) and Non-Responder (NR) mice. In Responders, tumor Cluc levels initially peaked but then declined, nearly disappearing by day 20. In contrast, Cluc levels in the NR group continued to rise. Based on these findings, we conducted a follow-up experiment to analyze brain activity during the first 20 days post-tumor implantation (**Figure 2a**) to explore the factors distinguishing the CR/PR group from the NR group. Following tumor implantation and iNG34 oHSV injection, we collected time-course data using MRI (**Figure 2b**) and MM-BLI (**Figure 2c**). All brains were harvested for formalin-fixed paraffin-embedded (FFPE) tissue analysis. MRI sequences, including T2-weighted imaging (T2WI) and T1-weighted imaging (T1WI) with and without contrast enhancer, were analyzed alongside Fluc, Rluc, and Cluc signals. Due to the short-term nature of the experiment, survival tracking was not feasible; therefore, we used MM-BLI and MRI T2LIV to categorize the mice as Responders or Non-Responders.

**Figure 2.**
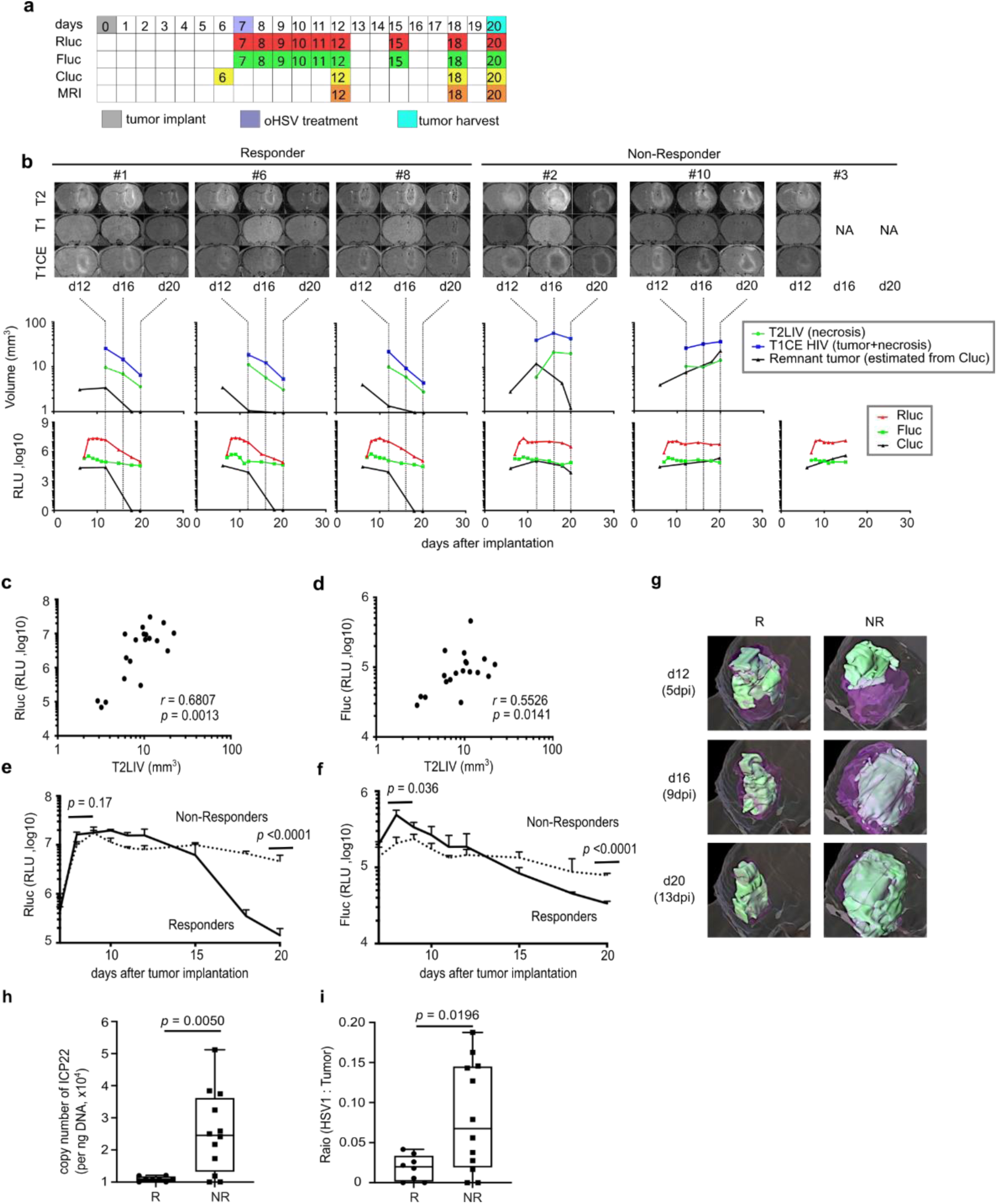
Short-term analyses correlating MRI with MM-BLI and iNG34 oHSV genome copy numbers. **a,** Scheme of scheduling MM-BLI and MRI, and procedures. **b,** (**upper**) time-series of MRI images of individual mouse after U87ΔEGFR-Cluc implantation. **(center)** Volumetric analyses of T2LIV and T1 hyper-intense volume (HIV) with estimated tumor volumes from serum Cluc. (**bottom**) MM-BLI longitudinal assays are shown below. Timelines link the MRI sequences, volume calculations, and MM-BLI data. Based on MRI and MM-BLI results, mice were classified as Responders (#1, 6, 8) or Non-Responders (# 2, 3, 10). Note that mouse #3 died at day 13. **c, d,** Correlation plots between T2LIV volume (representing iNG34 oHSV cytotoxicity and necrosis) and iNG34 oHSV Rluc (**c**) and Fluc (**d**) activity. Spearman’s rank correlation coefficients and p-values are shown. **e,f,** Temporal analysis of iNG34 biomarkers (**left;** Rluc, **right;** Fluc). The bar graph shows average maximum signals, and the slope of signal changes over time, analyzed using t-tests and linear mixed regression models, respectively. Mean + SEM. **h, i,** The box and whisker plots represent the quartiles of (**h**) iNG34 oHSV genome copy numbers (ICP22) and (**i**) the ratio of iNG34 biodistribution, measured by oHSV IHC, in day 20 brain FFPE specimens at the euthanasia timepoint for responder (R) and non-responder (NR) mice. **g,**

In the Responder group, a reduction in tumor T2 signal (correlated with T2LIV) and contrast-enhanced T1WI signals was observed, aligning with a decrease in Cluc expression. In contrast, the Non-Responder group showed no reduction in T2LIV or T1 contrast signal, corresponding to sustained Cluc levels. Both groups initially exhibited peaks in iNG34 oHSV Rluc and Fluc signals, with T2LIV positively correlating with Rluc (**Figure 2c**) and Fluc (**Figure 2d**) intensity. Responder mice had a significantly higher Fluc peak at 1-day post-injection (dpi) compared to Non-Responders (*p* = 0.036), followed by a faster decline in both Rluc and Fluc signals (*p* < 0.0001 for both) (**Figure 2e, f**). Furthermore, temporal analysis using contrast-enhanced T1WI for tumor progression and T2LIV for viral distribution revealed that while broader viral spread correlated with increased tumor destruction at 5 days post-infection, Non-Responders failed to eliminate tumors at 9-and 13-day post-infection, despite wider viral distribution and heightened activity (**Figure 2g**). Quantitative analysis of HSV DNA genomes (**Figure 2h**) and viral biodistribution (**Figure 2i**) in the FFPE tissue samples confirmed the persistent viral activity observed through MM-BLI in Non-Responders up to day 20. These findings underscore the key difference between the Responder and Non-Responder groups: Responders demonstrated rapid tumor lysis by iNG34, while in Non-Responders, where this initial robust response was absent, viral infection and replication continued within expanding tumor areas. Consequently, these tumors became increasingly resistant to HSV lysis over time.

### Correlation Between iNG34 MM-BLI-Assessed Infection/Replication, Tumor Viability, and Enhanced Biodistribution in Early-Stage Treated Tumors

Our previous observations indicated that early time points following iNG34 injection are crucial for distinguishing Responders from Non-Responders. To investigate this further, we analyzed brain tissues 2 days post-injection (dpi) of iNG34 (day 9 post-tumor implantation) (**Extended Data Fig. 4a**). IHC results showed that brains with peaks in Rluc expression at 2 dpi had widespread oHSV distribution throughout tumor areas, which correlated with slower tumor growth, as reflected by MM-BLI Cluc changes (**Extended Data Fig. 4b-e**). Some mice displayed increased Fluc signals, indicating potential Responders, while others with elevated Cluc levels and absence of Rluc and Fluc signal peaks at 2 dpi were classified as Non-Responders (**Extended Data Fig. 4c, e**). A significant correlation between reduced tumor growth and oHSV biodistribution (**Extended Data Fig. 4f**) suggests the predictive value of early iNG34 infection for therapeutic responsiveness.

### Viral and host transcriptomic changes associated with oHSV responsiveness

GBM is characterized by a tumor microenvironment rich in microglia and myeloid cells^22, 23^ . Using FFPE tissue sections (**Figure 3a**), we observed denser recruitment of Iba-1-positive microglia in areas of active iNG34 oHSV replication, marked by ICP4 expression during the early phase of virus administration (**Figure 3b**). In the Responder group, HSV-positive cells were detected across nearly the entire tumor area, along with CD45-positive immune cells. Both Responder and Non-Responder sections showed enrichment of Iba-1-positive microglia and F4/80-positive macrophages at the tumor-parenchymal border. However, fewer Iba-1-positive cells were found in distant parenchymal regions, suggesting reduced microglial and myeloid cell migration toward the tumor (**Figure 3c**). These findings indicate significant changes in the migration and activation status of brain-resident microglia and myeloid-derived immune cells in response to viral infection.

**Figure 3.**
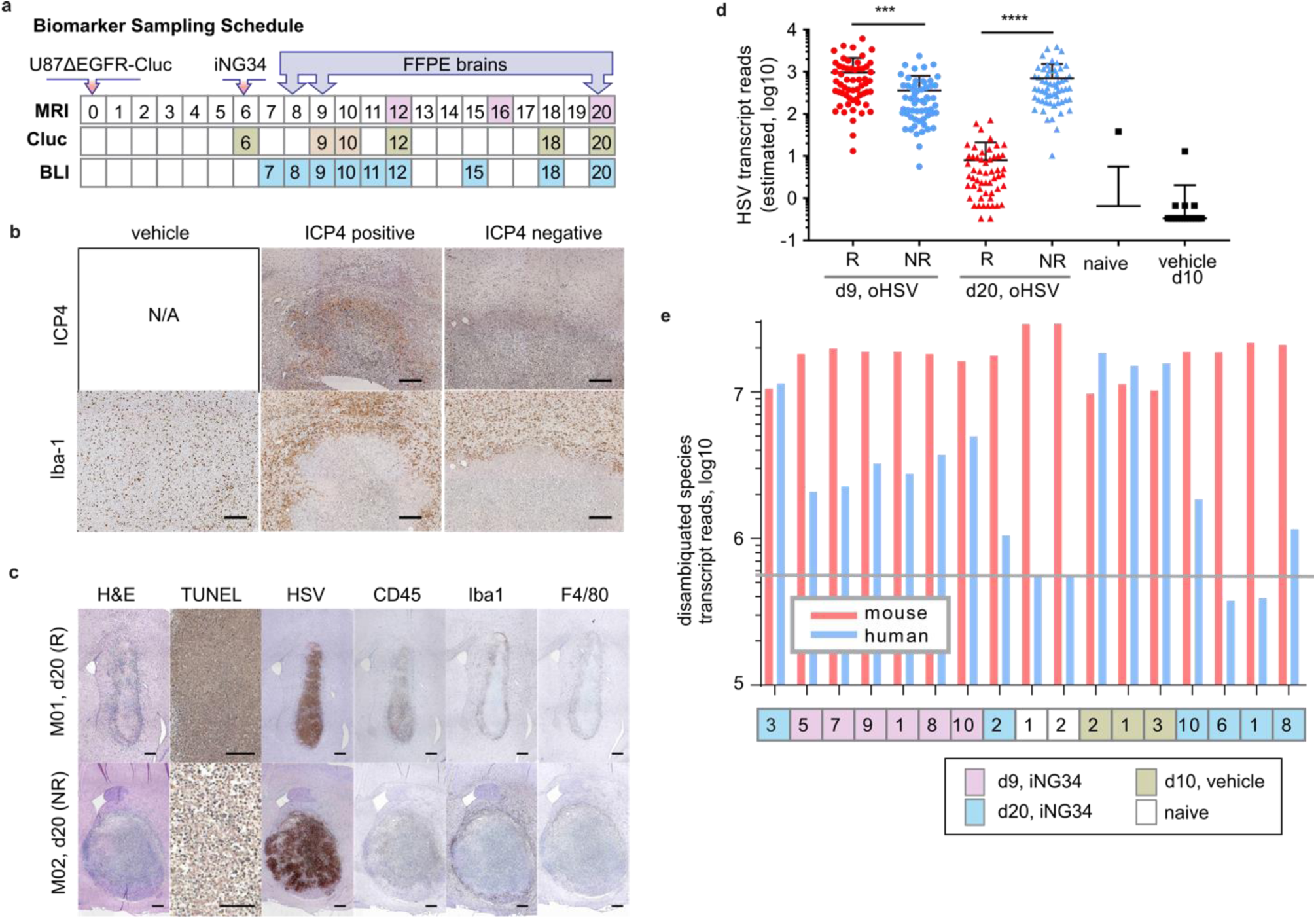
Multifactorial analyses validating biomarkers as surrogates for viral activities and investigating determinants of therapeutic response. **a,** Scheme of scheduling MM-BLI and MRI, and procedures. **b,** HSV-1 ICP4 and Iba-1 IHC stains for adjacent FFPE slices of identical mouse on the 3 dpi with iNG34. scale bar = 200 μm. **c**, Representative images of IHC of R and PR mouse brains on the 13 dpi with iNG34. scale = 200 μm for TUNEL, 100 μm for the rest. **d,** Scatter plots displaying the average expression of viral genes (58 genes in total) at 2 dpi and 13 dpi for both R (red dots) and NR (blue dots) groups. P values were calculated using the Kolmogorov-Smirnov test. **e,** Bar graph depicting the disambiguated human (blue) and mouse (red) sequence reads from the same grafted sample RNA-seq data, aligned to the hg38 and mm10 genomes, excluding ambiguous reads.

Our previous results from MM-BLI, HSV DNA analysis, and IHC indicated that the Responder group retained viral DNA and proteins, even though bioluminescent signals had dropped nearly to background levels (as seen in **Figure 2b, h, and 3c**). This finding suggests that while viral distribution remains confined within the tumor, it is no longer expanding. This observation correlates with the absence of viable tumor cells, as evidenced by undetectable Cluc in the serum.

To corroborate these findings, we performed bulk RNA-seq analysis on RNA extracted from FFPE specimens. The estimated number of HSV gene transcripts derived from sequence reads indicated higher viral activity in the Responder group at day 9, which became nearly undetectable by day 20 (**Figure 3d**). In contrast, the Non-Responder group exhibited persistent viral activity up to day 20.

We determined tumor viability based on human-derived transcripts, due to the xenograft model to differentiate between human tumor transcripts and mouse-derived transcripts (**Figure 3e**). Consistent with viral gene abundance, human transcript levels in the Responder group were significantly lower than in the Non-Responder group by day 20, reflecting tumor viability as indicated by Cluc levels. On the other hand, Responder and Non-Responder at early time point (day 9) show similar human transcript levels, yet lower than non-treatment group (vehicle), supporting overall therapeutic benefit of iNG34 oHSV in their survival course (**Figure 1a**).

These findings collectively demonstrate that multi-modality bioluminescence imaging (MM-BLI) effectively mirrors tumor viability and viral activity. Moreover, our results suggest that microglia and other myeloid cells actively migrate to areas of high viral activity, potentially influencing oHSV therapeutic responsiveness in later stages.

### PLX3397 treatment augments oHSV activity within tumor microenvironment

To investigate the effect of tumor-associated microglia ablation on OV-mediated tumor lysis, we employed PLX3397, a small molecule inhibitor targeting colony-stimulating factor-1 receptor and c-Kit (**Figure 4a**). We first confirmed that PLX3397 doesn’t affect oncolytic effects in *in vitro* cultured condition and doesn’t have toxicity in U87ΔEGFR-Cluc and G9-Cluc cells (**Extended Data Figure 5**). Using the U87ΔEGFR-Cluc orthotopic xenograft model, we observed that PLX3397 treatment alone had no significant impact on tumor growth or survival time (**Figure 4b**).

**Figure 4.**
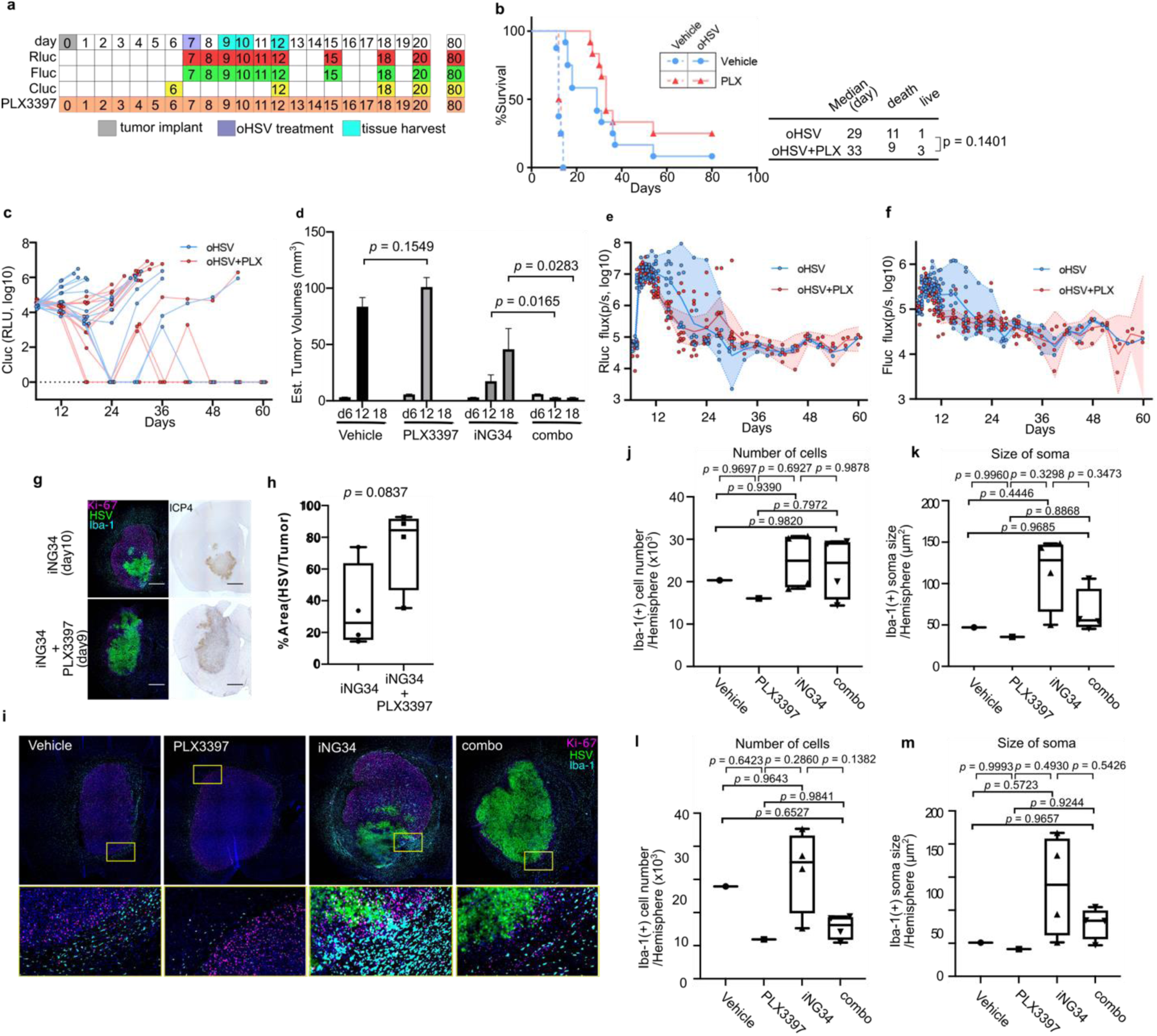
PLX3397 augments oHSV replication in tumor microenviroment by suppressing microglia recruitment to the infected area. **a,** Experimental timeline for MM-BLI (Rluc, Fluc, and Cluc) and PLX3397 treatment schedules following U87ΔEGFR-Cluc implantation, with n=8 (Vehicle with or without PLX3397) and n=12 (iNG34 with or without PLX3397). MM-BLI (Rluc; red, Fluc; green, Cluc; yellow box) and tissue harvests (light blue) are shown. Rluc and Fluc assays were conducted every 3 days after day 20, while Cluc assays were performed every 6 days after day 20, with all mice euthanized at day 80 for tumor analyses. **b,** Kaplan-Meier survival comparison for U87ΔEGFR-Cluc i.c. xenografts; p-values from the Log-rank test. **c,** Temporal changes in serum Cluc levels for individual mice in iNG34 (oHSV) with or without PLX3397. **d,** Volumetric analysis of tumors estimated from serum Cluc levels, with unpaired t -test results shown. **e, f,** MM-BLI results (**e;** Rluc, **f;** Fluc) presented as scatter plots (dots), non-linear regression curves (solid lines), and two-sided 95% confidence intervals (shaded areas). **g,** Representative images of multiplex immunofluorescence (IF) (left; HSV1; green, Iba-1; cyan, Ki-67; purple, DAPI; blue) and ICP4 IHC (right) at 10 dpi (iNG34 alone) and 9 dpi (iNG34 + PLX3397); scale bar = 1 mm. **h,** Box-and-whisker plot showing the percentage of virus-infected areas in U87ΔEGFR-Cluc i.c. tumors during the acute phase of virus infection (days 9-10); unpaired t-test results are shown. **i,** Representative images of multiplex IF (left; HSV1; green, Iba-1; cyan, Ki-67; purple, DAPI; blue). **j-m,** Box-and-whisker plots of the number of Iba-1+ cells (**j**; in hemispheres, **l**; in residual tumors) and soma size (**k**; in hemispheres, **m**; in residual tumors) in U87ΔEGFR-Cluc i.c. tumors during the acute phase of virus infection (days 9, 10, and 12 for iNG34 or iNG34 + PLX3397, day 13 for control (HBSS) or PLX3397). Data are represented as individual values, with N=1 for control (HBSS) or PLX3397, and N=4 for iNG34 or iNG34 + PLX3397. Tukey’s multiple comparisons tests are shown in the panels.

When combined with iNG34, PLX3397 altered survival curves compared to iNG34 monotherapy, though this change did not reach statistical significance (*p* = 0.1401). However, MM-BLI data revealed more promising outcomes: the combination therapy demonstrated improved viral biomarker patterns and notably suppressed tumor growth by day 18 (**Figure 4c-f**). These findings suggest that PLX3397 may modify the tumor microenvironment, rendering it more permissive to viral replication and oncolysis. We conducted quantitative multiplex immunohistochemistry (IHC) on brain samples. These samples were obtained 2-or 3-days post-virus inoculation in the U87ΔEGFR-Cluc brain tumor model. The primary aim of this analysis was to elucidate the status of Iba-1 positive cells (a marker for microglia and macrophages) and viral distribution *in vivo* (**Figure 4g, h**). Notably, the combined administration of PLX3397 resulted in an increased distribution of oHSV (**Figure. 4i**). The increase of the number and soma size of Iba-1 positive cells after virus inoculation, indicative of activation of microglia^24^, was hampered by PLX3397 (**Figure 4j-m**).

Overall, we demonstrated that MM-BLI effectively tracks tumor viability and viral activity in a glioblastoma model treated with oHSV, showing that early viral infection and replication are crucial for the therapeutic response. Our findings suggest that early viral dynamics and the tumor microenvironment, particularly the role of microglia, are key determinants of success, with the combination of PLX3397 emerging as an effective strategy to enhance oHSV treatment.

## Discussion

In this study, we hypothesized that integrating multiple imaging modalities, combined with the use of different luciferases to track various stages of OV infection/replication and tumor viability, would enhance our understanding of the factors critical for OV therapy success. We demonstrated that: (1) MM-BLI enables longitudinal, in vivo measurement of the kinetics of OV infection/replication and tumor viability; (2) MM-BLI correlates with OV anatomical distribution as measured by T2LIV MRI, oHSV IHC, and oHSV genome copy numbers; (3) MM-BLI patterns can be used to assess survival outcomes following OV therapy in a mouse model of human GBM; and (4) multimodal integration of MM-BLI, MRI T2LIV, and OV tumor transcriptomic analysis reveals that therapeutic success in this mouse model significantly correlates with the early phases of OV infection and replication in the injected tumors. Our findings support that the early stages of OV infection and replication are critical for determining therapeutic outcomes, underscoring the importance of early viral dynamics in the success of oncolytic virotherapy. Furthermore, the congregation of microglia at the infection site was discovered as a critical determinant of infection efficiency during this phase. Notably, ablating microglia with PLX3397significantly improved infection efficiency during the acute phase, leading to survival benefit.

We validated various modalities for analyzing oHSV tumor biodistribution, including T2WI of MRI, oHSV IHC, genome copy numbers, and MM-BLI. We demonstrated that early peaks in oHSV infection (Rluc) and replication (Fluc) within 2 dpi were significantly higher in responder mice, correlating with improved oHSV biodistribution, more genome copies, and reduced tumor growth. At later phases (13 dpi), Non-Responders showed increased infection and replication, but this did not improve survival outcomes. These findings suggest that the early phase of oHSV infection and replication is critical for positive outcomes, while later attempts to control rapidly growing tumors are less effective. Our recent clinical trial of CAN3110 (previously known as rQNestin34.5v2) also shows that persistent HSV antigens in patient tumors seem not to be beneficial to survival. Enhancing virotherapy may require technical, engineering, or pharmacological improvements focused on the early stages of oHSV infection. This raises the hypothesis that the effectiveness of oHSV therapy might be dependent on the rapid activation of the immune response during the initial infection phase, which could prevent tumor progression before resistance mechanisms develop. Further investigation into the molecular mechanisms governing this early-phase efficacy, particularly the interactions between viral replication and the tumor microenvironment, is essential to guide the development of optimized therapies. Understanding these processes will be crucial for designing interventions that either enhance early viral activity or mitigate the factors contributing to reduced efficacy in later stages of tumor growth.

The key advantage of MM-BLI is that it requires fewer resources and labor, allowing for longitudinal monitoring of oHSV and tumor kinetics in the same mouse, something not achievable with MRI or IHC/genome analysis alone. It can serve as a screening tool for testing different oHSV delivery methods (e.g., CED^28, 29^), engineered oHSVs with better biodistribution^30, 31^, and various combinations with therapeutic agents^32^. However, there are limitations. In immunocompetent syngeneic mouse GBM models, humoral immune responses against Cluc rise at later stages, making tumor volume measurement unreliable. Hemorrhages in treated tumors also interfered with light detection, making Rluc and Fluc values less interpretable. Additionally, light scattering *in vivo* can affect signal intensity based on the distance of the emission source from the detector^33^. Intratumoral necrosis can further complicate luciferase expression and light detection. Thus, while MM-BLI is a useful screening tool, integrating it with other assay modalities provides the most comprehensive approach for analyzing virotherapy.

In conclusion, we demonstrated that MM-BLI enables real-time tracking of OV-tumor dynamics, revealing that early-phase oHSV infection and replication are crucial for virotherapy success, with early peaks in viral activity closely correlating with improved outcomes, while later-phase infection does not enhance survival. Our findings also highlight the crucial role of microglia in oHSV infection efficiency with their removal promoting viral replication and tumor destruction. The integration of MM-BLI with other imaging and genomic techniques offers a promising avenue for a deeper understanding of virotherapy and the development of more effective treatments.

## Methods

### Cell lines

Vero (African green monkey kidney), U87ΔEGFR (human glioma), G9 (patient-derived glioma) were thawed from frozen stocks and have been previously utilized in the lab. They were cultured in DMEM (Thermo Fisher Scientific) supplemented with 10% fetal bovine serum (FBS, Sigma-Aldrich) and 100 U/mL penicillin-streptomycin (Thermo Fisher Scientific). U87ΔEGFR-Cluc and G9-Cluc cell lines were generated from U87ΔEGFR and G9, respectively, by stable transduction with a lentiviral vector expressing Cluc and a puromycin resistant gene. Cluc-expressing cells were selected by puromycin. Correlation of Cluc in cultured media with the number of cells seeded in 6-well plates at 10^2^, 10^3^, 10^4^, or 10^5^ cells/well were analyzed in U87ΔEGFR-Cluc and G9-Cluc cell lines. Cluc expression was confirmed and Pearson’s correlation coefficients with five biological replicates are shown in Extended Data Figure 2.

### Engineering of iNG34

iNG34 was generated using the NG34 backbone^15^, as described in Extended Data Figure 1a. Briefly, a BamHI -NotI fragment containing the Green Renilla (Gr) luciferase cDNA from the commercially available PTK-Green Renilla Luc (Pierce) plasmid was cloned into the pT-oriSIE4/5 plasmid DNA at the EcoRV site^25–27^, to generate IE4/5-Grluc. Then, the BstBI-BbsI fragment of IE4/5-Grluc was inserted into pT-NG34-gCRliFluc to generate the shuttle DNA vector for the fHSVQuik1 cloning system, as previously described ^25, 27^. Correct insertion of the fragment was confirmed by pulsed field gel electrophoresis (PFGE).

### Bioluminescence imaging *in vitro*

For temperature shift experiments, Vero cells were plated in 6-well plates at a concentration of 2×10^5^ cells/well. A day later, the cells were exposed to 3×10^5^ pfu of iNG34. After 1 hour incubation in 4℃, cells were washed to remove iNG34 and 2 mL of DMEM supplemented with 2% FBS, preheated to 37 or 40℃, was added to the plates in a CO_2_ incubator at 37 or 40℃, respectively. Rluc or Fluc signal luminescence was measured 2, 4, 6, 8, 14, 20 hours after infection using the instructions provided by the commercially available kits for *Renilla* Luciferase Assay System (Promega) or Luciferase Assay System (Promega) respectively: bioluminescence emission at 520 nm and 590 nm, respectively, was assayed using the POLARstar Omega Plate Reader Spectrophotometer (BMG Labtech) and reported as relative fluorescence unit (RLU).

For inhibition assays with PAA, U87ΔEGFR-Cluc cells were plated in 6-well plates at a concentration of 3×10^5^ cells/well. A day later, the cells were exposed to 3×10^4^ pfu of iNG34. After 1 hour incubation in 37℃, cells were washed to remove iNG34 and 2 mL of DMEM supplemented with 2% FBS with or without PAA (0.4mg/mL) was added to the plates in a CO_2_ incubator at 37℃. Rluc or Fluc signal luminescence was measured 24, 48, 72 hours after infection as described above.

For Cluc assays, 20 μL of cell supernatants were assayed following the instructions in the Pierce *Cypridina* Luciferase Flash Assay Kit (16169, ThermoFisher Scientific) and measured in the POLARstar Omega Plate Reader Spectrophotometer (BMG Labtech). Signal was measured 1 second after substrate injection and quantified as RLU.

### Bioluminescence imaging *in vivo*

For *in vivo* Fluc bioluminescent imaging, D-luciferin (Promega; dissolved in sterile D-PBS) at a dose of 3 mg per 20 g body weight was injected intraperitoneally. For Rluc bioluminescent imaging, Coelenterazine (Nanolight Technology; dissolved in sterile water) at a dose of 50 mg per 20 g body weight was injected intravenously. All procedures were carried out under anesthesia with isoflurane vaporization. Light-emission was acquired with the IVIS Lumina LT with Living Image^®^ software (Perkin-Elmer) every 60 seconds. Peak signal intensities with Cy5.5 or DsRed emission filters, respectively, were measured as total flux (photons/sec). For the Cluc assay, 20 μL of 1/30 diluted (U87ΔEGFR-Cluc) serum were assayed using instructions provided with the Pierce *Cypridina* Luciferase Flash Assay Kit (16169, ThermoFisher Scientific) and measured in a POLARstar Omega Plate Reader Spectrophotometer (BMG Labtech). Signal emission was measured 1 second after substrate injection and quantified as RLU.

### Animal experiments

Six- to 8-week-old female athymic nude mice were purchased from Envigo. All experimental procedures using animals were carried out under an animal protocol reviewed and approved by the BWH’s IACUCs and performed in accordance with relevant guidelines and regulations. Dissociated U87ΔEGFR-Cluc or G9-Cluc tumor cells (100,000 or 50,000 cells in 2 μL of Hanks’ buffered saline solution, HBSS, respectively) were injected stereotactically with a Kopf stereotactic frame (David Kopf Instruments) into the right frontal lobe of mice (ventral 3.0 mm, rostral 0.5 mm, and right lateral 2.0 mm from bregma). Six days after implantation, tumor load was calculated based on the Cluc levels in serum as described above and then mice were assigned to different groups for treatment (iNG34 (2×10^5^ pfu) or HBSS), ensuring equivalence in tumor load for either group. Bioluminescence imaging for Rluc and Fluc was performed the first 6 consecutive days starting from 4-6 hours after virus inoculation on day 7, followed every 3 days after that. Tumor load was also monitored by Cluc assay every 6 days starting from day 6. For immunohistochemistry and extraction of DNA and RNA, mice were sacrificed at the designated time points, and brains were collected after perfusion with PBS and 10% neutral buffered formalin solution (Sigma-Aldrich).

### Diet Preparation and PLX3397 Administration

PLX3397 was purchased from MedChemExpress and incorporated into the standard chow diet at a concentration of 290 mg/kg diet, ensuring uniform distribution of the compound by Research Diets, Inc. Control animals were fed the same diet without PLX3397.

### Magnetic Resonance Imaging (MRI)

Intracranial tumors were imaged on a 3T MRI (Bruker Biospin) under anesthesia with an isoflurane vaporizer. First, T2WI were acquired using the following parameters: repetition time (TR) = 4,993.715 msec; echo time (TE) = 66.54 msec; 20 slices; thickness = 0.5 mm; field of view (FOV) = 20 x 20 mm^2^; matrix = 192 x 192; number of averages (NA) = 4. Then, T1WI without and with contrast (100 μL 10-fold diluted gadopentetate dimeglumine) were acquired with the same geometry using the following parameters: TR = 1,000 msec; TE = 10 msec; 20 slices; thickness = 0.5 mm; field of view (FOV) = 20 x 20 mm^2^; matrix = 192 x 192; NA = 3.

### Tumor volume measurements from assays of Cluc *in vitro* and *in vivo*

To validate the quantitative correlation between tumor cell number and Cluc level in supernatant, 100, 1,000, 10,000, 100,000 cells of U87ΔEGFR-Cluc or G9-Cluc cells were seeded onto 24-well plates with 1mL of DMEM supplemented with 10%FBS in triplicate and incubated at 37℃ in an atmosphere containing 5%CO_2_. Twenty μL of supernatant were used for Cluc assays, as described above. To estimate the tumor volumes from Cluc levels in serum, 50,000 cells of U87ΔEGFR-Cluc cells or G9-Cluc cells were stereotactically implanted in the right frontal lobes of athymic mice and T2WI were acquired on day 6, 12 and 14 for U87ΔEGFR-Cluc, on day 8, 14 and 16 for G9-Cluc, respectively, as described above. Sera were also collected on the same day as MRI acquisition and Cluc levels were measured as described above. Tumor volumes were calculated by a manual segmentation of tumors using the Horos open-source medical imaging viewer (Horos.org; https://horosproject.org/)

### Extraction of DNA and RNA from FFPE samples

Total DNA and RNA were extracted from Formalin fixed paraffin embedded (FFPE) from sections immediately adjacent to the sections used for immunohistochemical staining, using instructions from the commercially available QIAamp DNA FFPE Tissue Kit (Qiagen) and RNeasy FFPE Kit (Qiagen), respectively. Purified RNA was reverse-transcribed into cDNA using instructions from the commercially available iScript cDNA Synthesis Kit (Bio-Rad).

### Quantitative PCR

The following primers and probes were synthesized by Thermo Fisher Scientific and used throughout the study: ICP22 For: GAGCAGCGCACGATGGA; ICP22 Rev: CCCGAAACAGCTGATTGATACA; ICP22 Probe: 6FAM-CCG ACC TGG GCT ACA TGC GCC-TAMRA; gC For: AGGTCCTGACGAACATCACC; gC Rev: GCCCGGTGACAGAATACAAC; gC Probe: VIC-GGGACTCCTGGTGTACGACA-QSY. Taqman Universal PCR Master Mix (Thermo Fisher Scientific) and Applied Biosystems 7500 Fast Real-Time PCR system (Thermo Fisher Scientific) were used according to the manufacturer’s protocol. The amplification step consisted of 40 cycles at 95℃ for 20 seconds followed by 60℃ for 20 seconds. Serial tenfold dilutions of were used as templates to generate standard curves. oHSV genomic DNA was calculated and measured as copy numbers per ng DNA or RNA.

### Immunohistochemistry

Mouse brains were fixed in 10% neutral buffered formalin solution (Sigma-Aldrich) overnight and embedded in paraffin. Serial sections were sectioned to a thickness of 4 μm with a microtome. Tumors sections were analyzed for HSV-1, ICP4, CD45, Iba1, F4/80 and HIF-1α as follows: tissue slides were deparaffinized and rehydrated using xylene and gradually decreasing concentrations of ethanol. Endogenous protein and endogenous peroxidase were blocked using 1.5% normal goat serum (S-1000; Vector Laboratories) and with 0.3% H_2_0_2_/methanol, respectively. Antigen retrieval was performed using a citrate buffer at a pH 6.0 and with microwave heating at 95℃ for 20 minutes. Sections were incubated at room temperature for 1 hour with a primary antibodies: HSV (1:2000, B0114 (Dako)), CD45 (1:25, 553076 (BD)), Iba1 (1:500, ab178846 (abcam)), F4/80 (1:2000, ab6640 (abcam)), HIF-1α (1:200, ab51608 (abcam)) After a 30-min incubation with Goat Anti-Rabbit IgG H&L (HRP polymer)(ab214880 (abcam)) or Anti-Rat IgG H&L (HRP polymer)(ab214882 (abcam)). A DAB Substrate Kit (SK-4100; Vector) was used for detection of secondary antibody staining. All sections were counterstained with hematoxylin for 2 min, dehydrated with gradually increasing concentrations of ethanol, and fixed with xylene.

TUNEL assay was performed using Click-iT™ TUNEL Colorimetric IHC Detection Kit (Thermo Fisher Scientific) according to the manufacturer’s instructions. **Quantification:** HSV positive areas and tumor areas were measured by manual segmentation with NIS-Elements Imaging Software (Nikon Instruments Inc.). Outline of the area to be quantified were traced by a freehand selection tool.

### Multiplex Immunofluorescence Staining

Multiplex immunofluorescence staining was performed using the Opal 4-Color IHC Kit (Akoya Biosciences) according to the manufacturer’s instructions. Primary antibodies specific for Ki-67, HSV-1 and Iba-1were sequentially applied, each followed by Opal polymer horseradish peroxidase (HRP) secondary antibody and tyramide signal amplification (TSA) using Opal dyes Opal690, Opal520, and Opal570, respectively. Nuclei were counterstained with DAPI for 5 minutes. Slides were mounted using ProLong Gold Antifade Mountant (Thermo Fisher Scientific) and stored in the dark at 4°C until imaging. Immunofluorescent images were captured using a DMi8 Widefield Microscope (Leica). Each fluorophore was imaged at its respective wavelength, and spectral unmixing was performed to separate overlapping signals. The resulting images were analyzed using Image J to quantify the number and the soma size of Iba-1 positive cells.

### RNA sequencing analysis

**Sample Preparation:** The samples were sectioned at 4 µm thickness using a microtome and de-paraffinized using xylene or other solvent. **RNA Extraction and Quality Control:** Total RNA extraction was performed using the Qiagen RNeasy FFPE Kit following the manufacturer’s protocol, incorporating DNase treatment to remove genomic DNA contamination. RNA quantity and integrity were assessed using a NanoDrop spectrophotometer. **Library Preparation and Sequencing:** RNA-seq libraries were prepared using the NEBNext Ultra II RNA Library Prep Kit for Illumina according to the manufacturer’s instructions. The library quality was validated using the Agilent Bioanalyzer, and quantified with the Qubit fluorometer. Sequencing was performed on the Illumina NovaSeq 6000, generating 150 bp paired-end reads. **Data Analysis:** FASTAQ data were processed in the bcbio-nextgen pipeline (HMS O2) with GRCm38v98 (mm10) and hg38 genome data, aligned using STAR (2.6.1d). Novel transcripts were assembled and merged into the known annotation with StringTie, and counts were estimated using Salmon (0.14.2). For HSV-specific analysis, bcbio-nextgen was run with the hisat2 aligner using GenBank x14112, with Salmon used for count estimation. Differential gene expression was analyzed with DESeq2 in R, and GO-PCA analysis was performed using Python.

### Ethical approval declarations

All mouse experiments were approved by the Institutional Animal Care and Use Committee (IACUC) at Brigham and Women’s hospital following guidelines set forth by the National Institutes of Health Guide for the Care of Use of Laboratory Animals as well as by the Association for Assessment and Accreditation of Laboratory Animal Care (AAALAC).

---

**Extended Data Figure 1.**
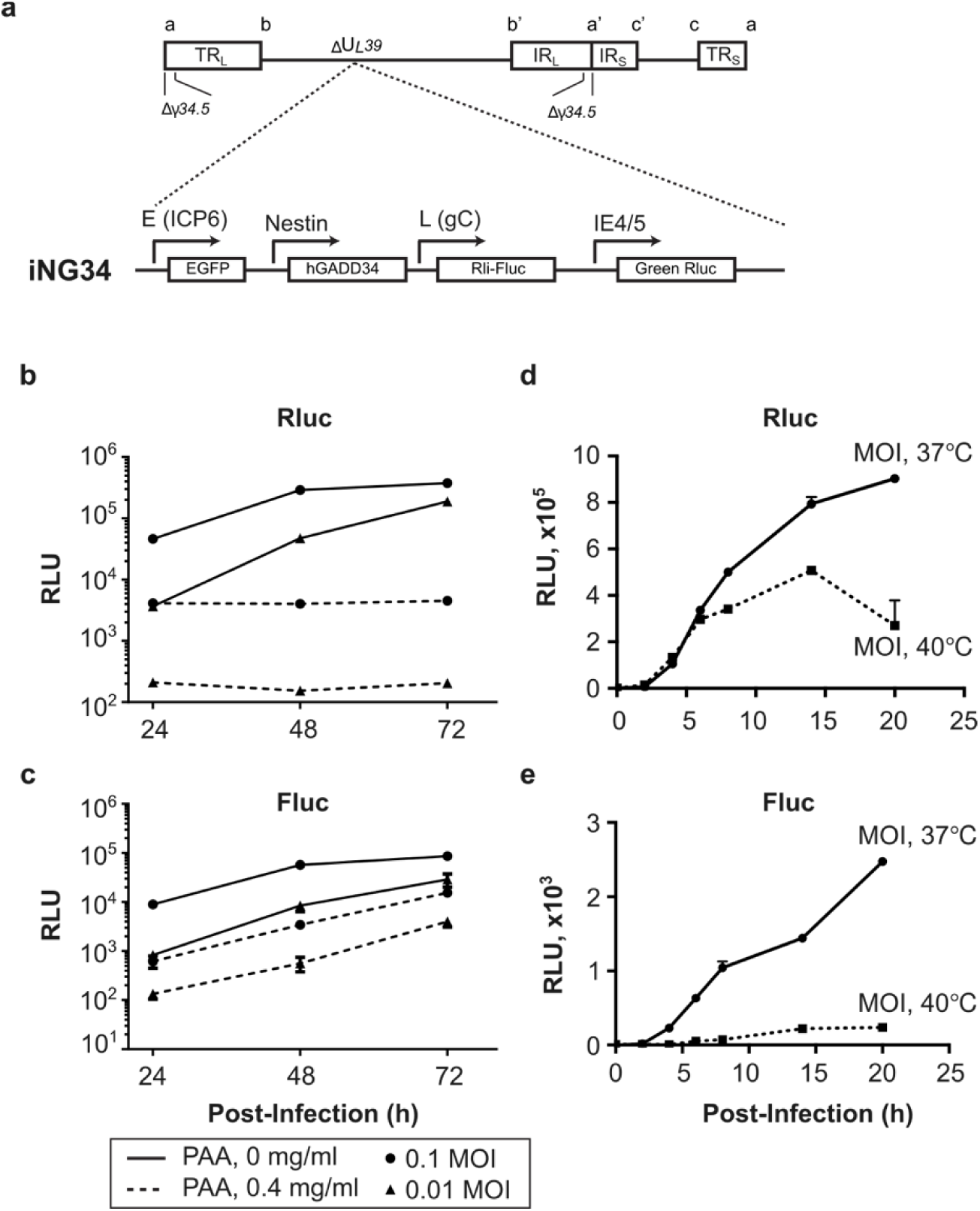
The structure of iNG34 and independence of each promoter for Rluc or Fluc. **a,** Schematic of the iNG34 construct showing viral immediate early (IE4/5) and late genes (gC) promoter-driven luciferase cDNAs (Rluc and Fluc) integrated in the UL39 locus with the NG34 backbone. **b-e,** Rluc (**b, d**) and Fluc (**c, e**) activities were assessed using lysates from U87ΔEGFR-Cluc cells (350,000 per well of 6-well plate) or Vero cells (200,000 per well of 6-well plate) infected with iNG34. Promoter activities were measured in the presence of PAA (0.4 mg/ml) (**b, c**) or at different temperatures (37 or 40 ℃) (**d, e**). Luminescence is expressed as Relative Light Units (RLU) on the y-axis. Mean ± SEM.

**Extended Data Fig. 2.**
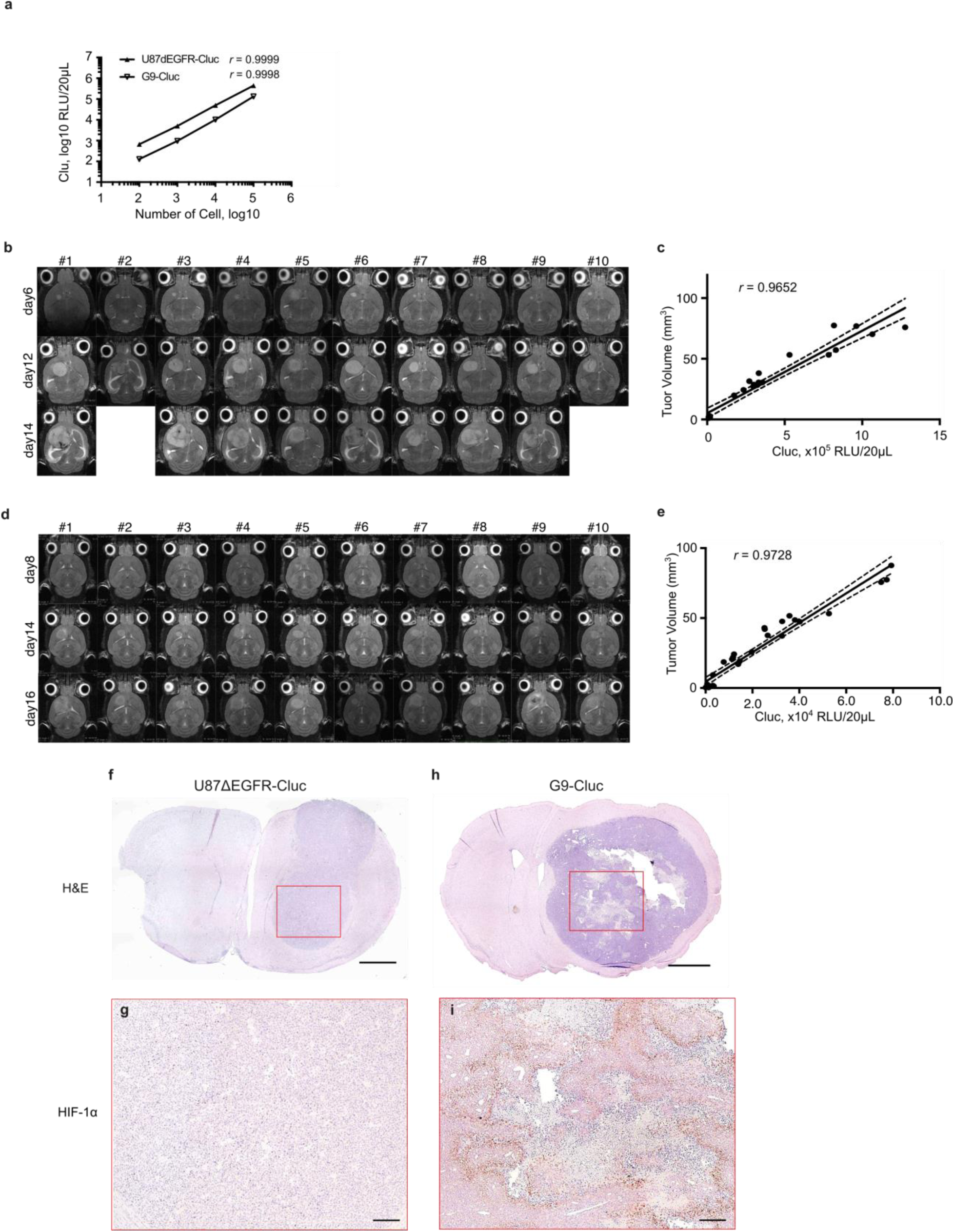
Validation of Cluc as a quantitative tumor biomarker in histologically distinct human GBM lines. **a,** Correlation of Cluc in cultured media with vial cell numbers at 24 hours after the indicated cell numbers seeded in 6-well plates in both U87ΔEGFR-Cluc and G9-Cluc cell lines. Pearson’s correlation coefficients with five biological replicates. Mean + SEM. **b-e,** Correlation between Cluc from serum and MRI volumetric measurements of brains from athymic mice bearing U87EGFR-Cluc (**b,c**) or G9-Cluc (**d,e**) intracranial tumors are shown. MRI T2-weighted images were taken on the indicated days post-implantation, and sera were collected at post-MRI. Pearson’s correlation coefficients are shown. **f-i,** Representative histology of U87EGFR-Cluc (**f,g**) and G9-Cluc (**h,i**) bearing brains harvested at day 10 and 22, respectively. Hematoxylin and eosin (H&E) stain (**f, h**) and immunohistochemistry staining with anti-HIF-1α antibody (**g, i**) are shown. Scales; 1 mm (f, h), 100 µm (g, i).

**Extended Data Figure 3.**
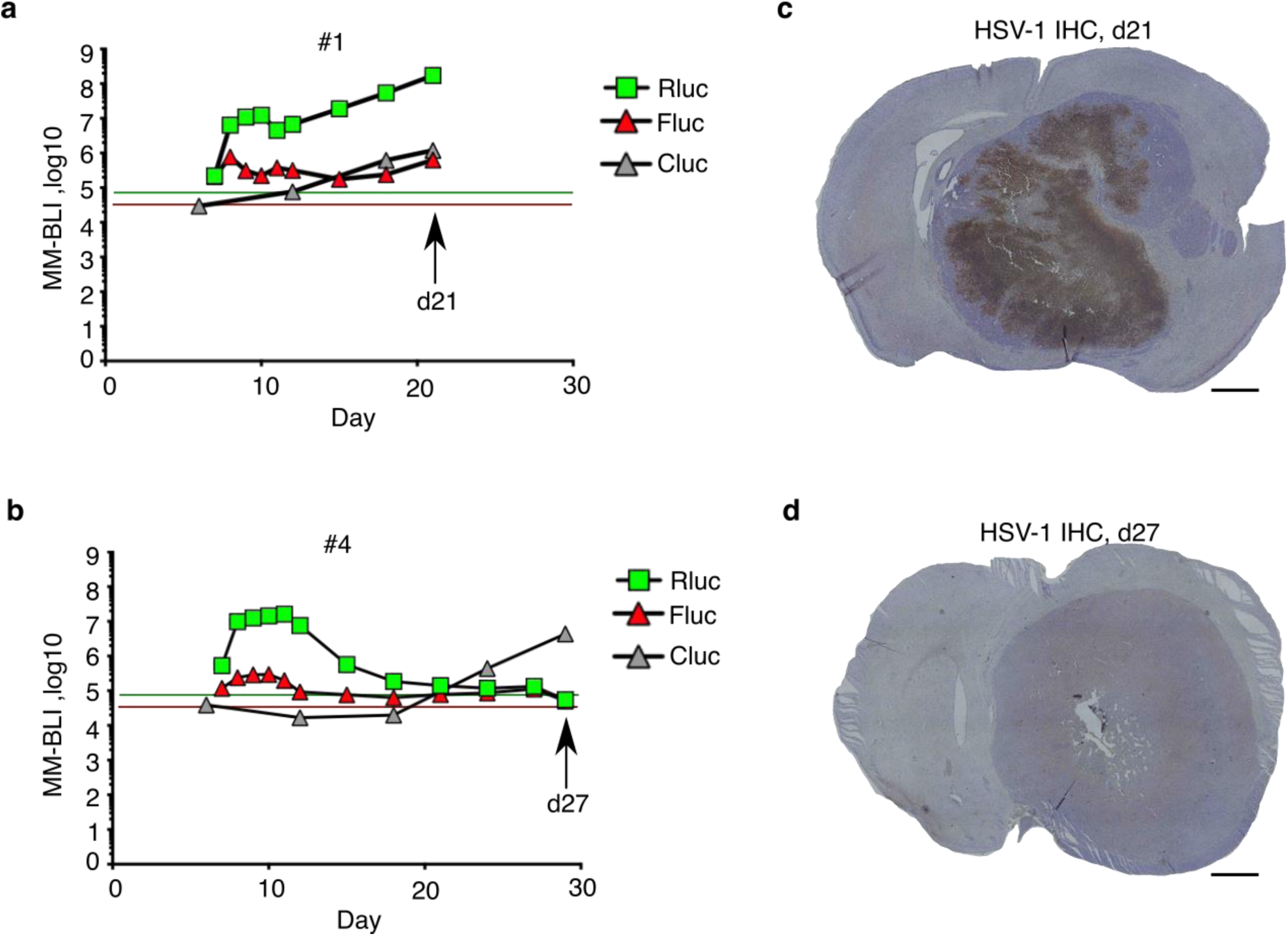
Two types of non-responder cases in iNG34-treated mice. **a-d,** U87ΔEGFR-Cluc tumors exhibit a wide viral distribution with cumulative MM-BLI (**a, c**) and a lack of remaining virus with decay following the peak of MM-BLI (**b, d**). FFPE brain tissues were collected at the times indicated by arrows in the plots. The y-axis shows flux of Rluc and Fluc as photons per second (p/s) or RLU per 10 µl serum, while the x-axis shows days after tumor implantation. Mouse individuals (#1, #6) correspond to those in Figure 1. Lines (green for Rluc, red for Fluc) indicate the baseline of background signals.

**Extended Data Figure 4.**
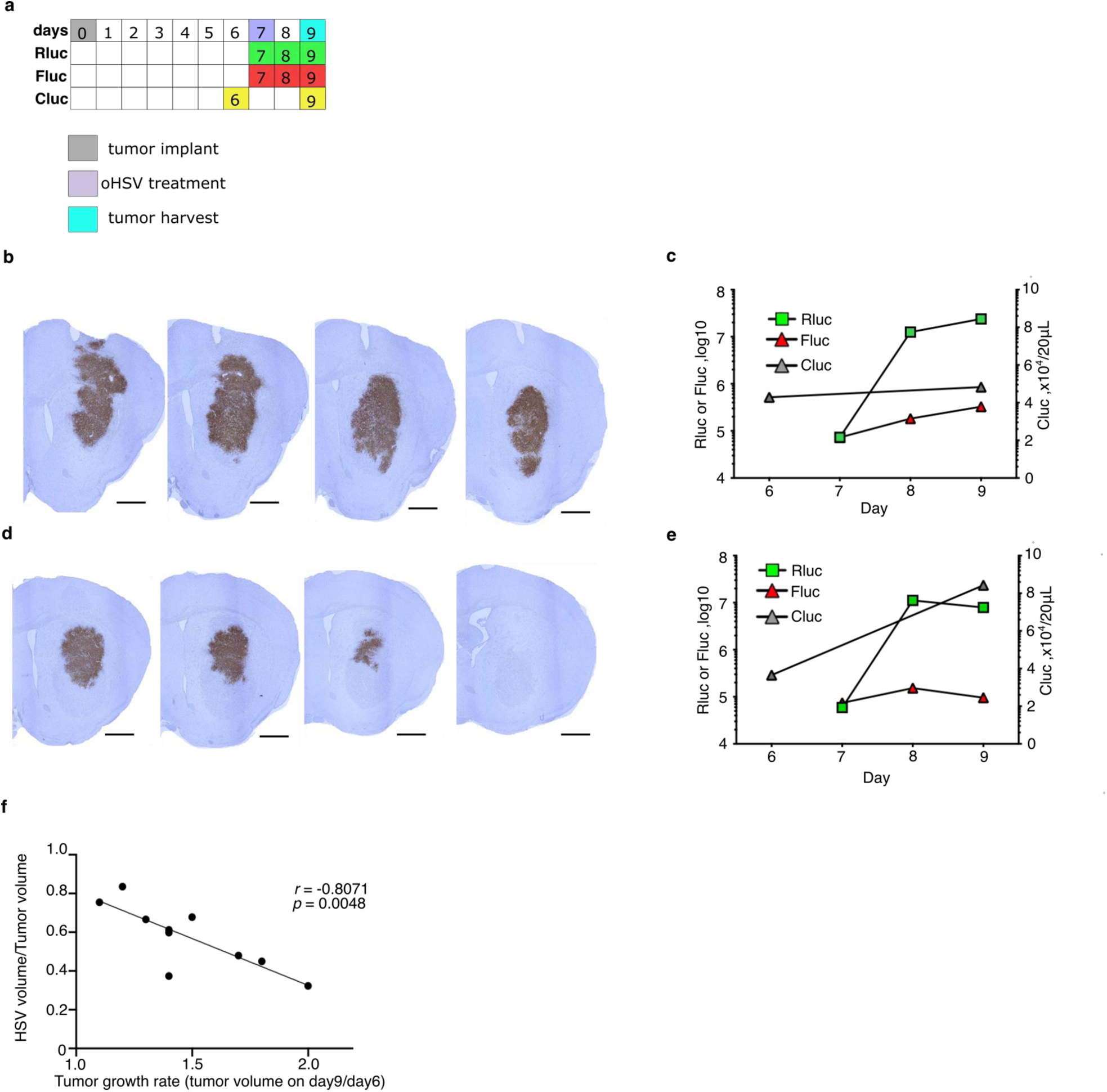
Short-term analyses correlating MM-BLI, IHC and tumor growth rates. **a,** Scheme outlining the scheduling of MM-BLI, tumor harvest, and associated procedures. **b, c,** Possible responder mouse: serial sections of HSV-1 IHC stains (**b**) and measurements of iNG34 oHSV Rluc, Fluc, and serum Cluc (**c**). d, e, Possible non-responder mouse: serial sections of HSV-1 IHC stains (d) and measurements of iNG34 oHSV Rluc, Fluc, and serum Cluc (**e**). Scale bar = 200 µm. **f,** Correlation plots showing the relationship between HSV volume/tumor volume (representing iNG34 oHSV biodistribution) and tumor growth rate (estimated tumor volumes from serum Cluc, ratio of day 9 to day 6), with Pearson’s rank correlation coefficient and p-value indicated.

**Extended Data Figure 5.**
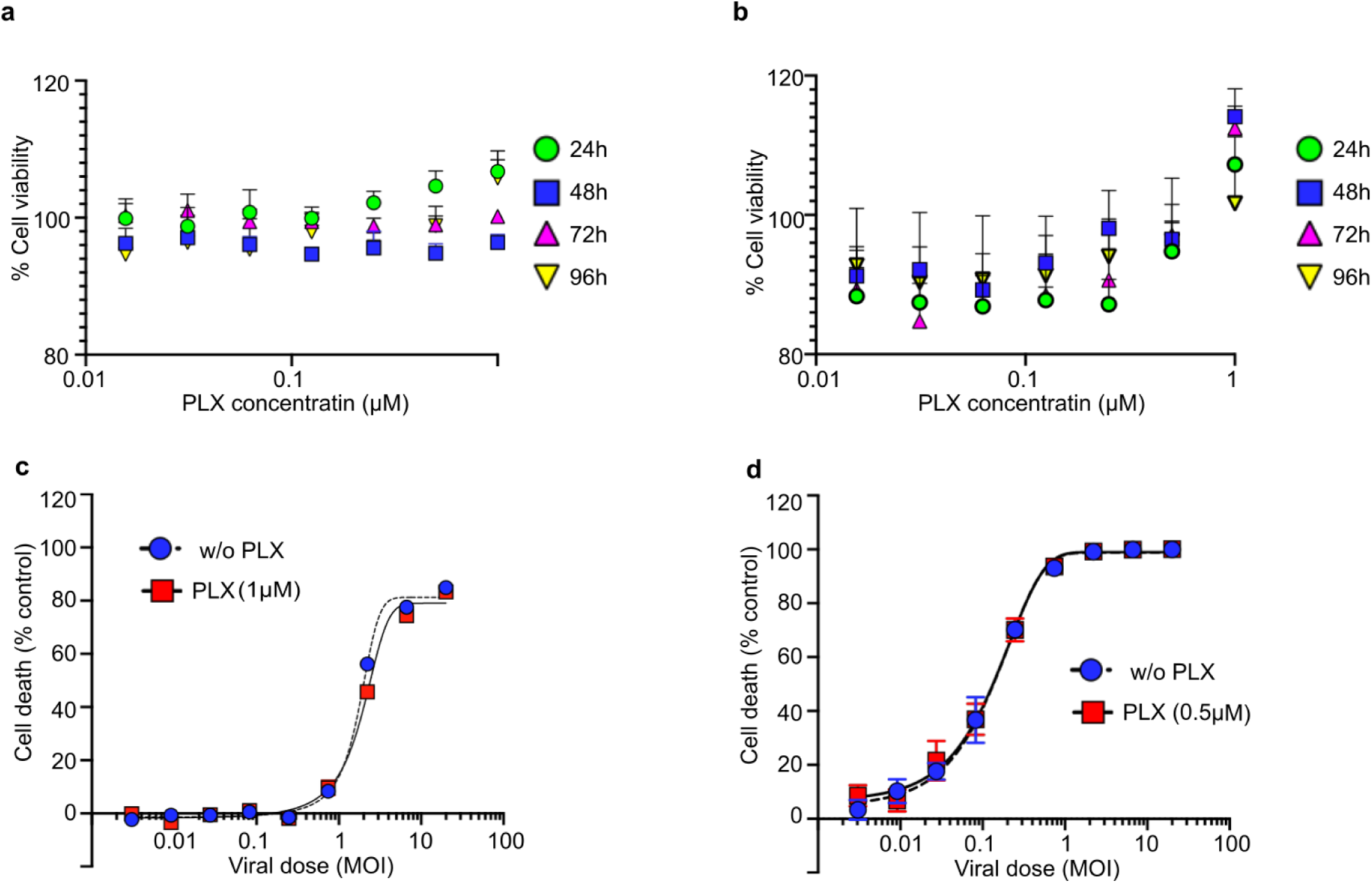
Effects of PLX3397 on tumor growth and viral activity in vitro. **a, b,** Cell proliferation assays show the impact of PLX3397 on U87ΔEGFR-Cluc (**a**) and G9-Cluc (**b**) cell viability over various concentrations and time points as indicated in plots. **c, d,** Cytotoxicity assays assess cell death in U87ΔEGFR-Cluc and G9-Cluc cells infected with iNG34 at different MOIs, with PLX3397 added at the concentrations shown in the plots. Symbols represent mean ± s.e.m. for cell viability and cell death from triplicate or sextuplicate cultures.

